# Versatile live attenuated SARS-CoV-2 vaccine platform applicable to variants induces protective immunity

**DOI:** 10.1101/2021.02.15.430863

**Authors:** Akiho Yoshida, Shinya Okamura, Shiho Torii, Sayuri Komatsu, Paola Miyazato, Shiori Ueno, Hidehiko Suzuki, Wataru Kamitani, Chikako Ono, Yoshiharu Matsuura, Shiro Takekawa, Koichi Yamanishi, Hirotaka Ebina

**Author notes:** These authors contributed equally (Akiho Yoshida, Shinya Okamura).

## Abstract

Live attenuated vaccines are generally highly effective. Here, we aimed to develop one against SARS-CoV-2, based on the identification of three types of temperature-sensitive (TS) strains with mutations in nonstructural proteins (nsp), impaired proliferation at 37-39°C, and the capacity to induce protective immunity in Syrian hamsters. To develop a live-attenuated vaccine, we generated a virus that combined all these TS-associated mutations (rTS-all), which showed a robust TS phenotype *in vitro* and high attenuation *in vivo*. The vaccine induced an effective cross-reactive immune response and protected hamsters against homologous or heterologous viral challenges. Importantly, rTS-all rarely reverted to the wild-type phenotype. By combining these mutations with an Omicron spike protein to construct a recombinant virus, protection against the Omicron strain was obtained. We show that immediate and effective live-attenuated vaccine candidates against SARS-CoV-2 variants may be developed using rTS-all as a backbone to incorporate the spike protein of the variants.

## INTRODUCTION

The COVID-19 pandemic, caused by the pathogen SARS-CoV-2, has had a serious impact on public health, with more than 500 million infection cases and over six million deaths reported worldwide (ourworldindata.org). To prevent the spread of COVID-19, several adenovirus-vectored and mRNA vaccines encoding the viral spike (S) protein gene have been developed and are presently widely used. These vaccines have been reported to induce robust humoral and cellular immune responses (Folegatti et al., 2020; Jackson et al., 2020; Sahin et al., 2021); however, there are concerns regarding adverse reactions caused by these vaccines, such as thrombosis and fever. Furthermore, variants of concern (VOC), which bear several mutations in the S protein, continually arise and contribute to the evasion of the humoral immunity generated against the ancestral S protein (Baum et al., 2020; Edara et al., 2021; Wibmer et al., 2021). To increase the induction of neutralizing antibodies against VOCs, such as the SARS-CoV-2 Omicron variant, most countries have been encouraging a third, and even a fourth, vaccine dose as a booster. Nevertheless, the antibody response induced by these vaccines is not persistent, demanding the development of alternative vaccines of different modalities to better control the ongoing pandemic.

Traditional live-attenuated vaccines are highly effective and have been successfully used against various diseases, including varicella, measles, and rubella viruses. These vaccines were developed by heterogeneous adaptation, isolating mutant viruses that cannot propagate in human cells (Makino et al., 1970; Parks et al., 2001; Sasaki, 1974; Shishido and Ohtawara, 1976; Takahashi et al., 1974; Zimmerman et al., 2018), or by isolating temperature-sensitive (TS) viruses that cannot replicate at the physiological human body temperature (Komase et al., 2006; Okamoto et al., 2016). In addition, a live influenza virus vaccine was developed by reassortment of viruses generated from a cold-adapted donor virus with temperature sensitivity-related mutations in six vRNA segments (Maassab and Bryant, 1999; Murphy and Coelingh, 2002). The attenuated phenotype of this virus was confirmed by its capacity to replicate at 25–33°C but not at 37°C (Cox and Dewhurst, 2015). Therefore, several groups have been developing live-attenuated vaccines against SARS-CoV-2. For example, Trimpert et al. and CODAGENIX are developing a codon-deoptimization strategy (Trimpert et al., 2021; Wang et al., 2021) and Seo et al. followed the cold-adaptation approach for isolating live-attenuated TS strains (Seo and Jang, 2020).

In this study, we obtained four TS SARS-CoV-2 strains from a clinical isolate by random mutagenesis. These strains exhibited low pathogenicity and protecting immunogenicity *in vivo*. We identified that mutations in *nsp3*, *nsp14,* and *nsp16* genes are associated with the TS phenotype, and generated a highly effective and safe live-attenuated vaccine candidate by combining these TS-related mutations. Moreover, this candidate vaccine could be adjusted with an appropriate S protein to generate vaccines for specific VOCs. We believe that this live-attenuated vaccine candidate is a promising platform to control the spread of the COVID-19 pandemic.

## RESULTS

### SARS-CoV-2 TS mutants show attenuated phenotype and induce protective immunity

To isolate the TS strains of SARS-CoV-2, we generated a library of viruses containing random mutations from the clinical isolate B-1 virus (accession number: LC603286) (Figure S1A). In total, 659 viral plaques were isolated from the library and screened. Vero cells were infected with all viral clones, cultured at 32 or 37°C, and monitored daily for cytopathic effects (CPE). During this process, we selected four TS strains that induced CPE at 32°C three days post-infection (dpi) but not at 37°C. To comprehensively analyze temperature sensitivity, we evaluated the growth kinetics of the isolated TS strains at 32, 34, and 37°C (Figures 1A and S1B). Under 32°C and 34°C culture conditions, all TS strains replicated comparably to the parent B-1 virus. However, at 37°C, the replication of all TS strains was relatively slower or smaller in scale than that of the parent strain. The growth of the H50-11 strain was delayed; however, its titer on day five was comparable to the viral titer peak of the B-1 virus on days two and three (p = 0.38). The L50-33 and L50-40 strains slightly proliferated at 37°C, and the viral titers were less than 10^4^ TCID_50_/mL, even at five dpi, and were 10^4^ times lower than the maximum titer of the B-1 strain. Interestingly, the A50-18 strain showed a unique phenotype, as no infectious viruses were detected in the culture medium at 37°C (the TCID_50_ was under the assay’s limit of detection).

**Figure 1.**
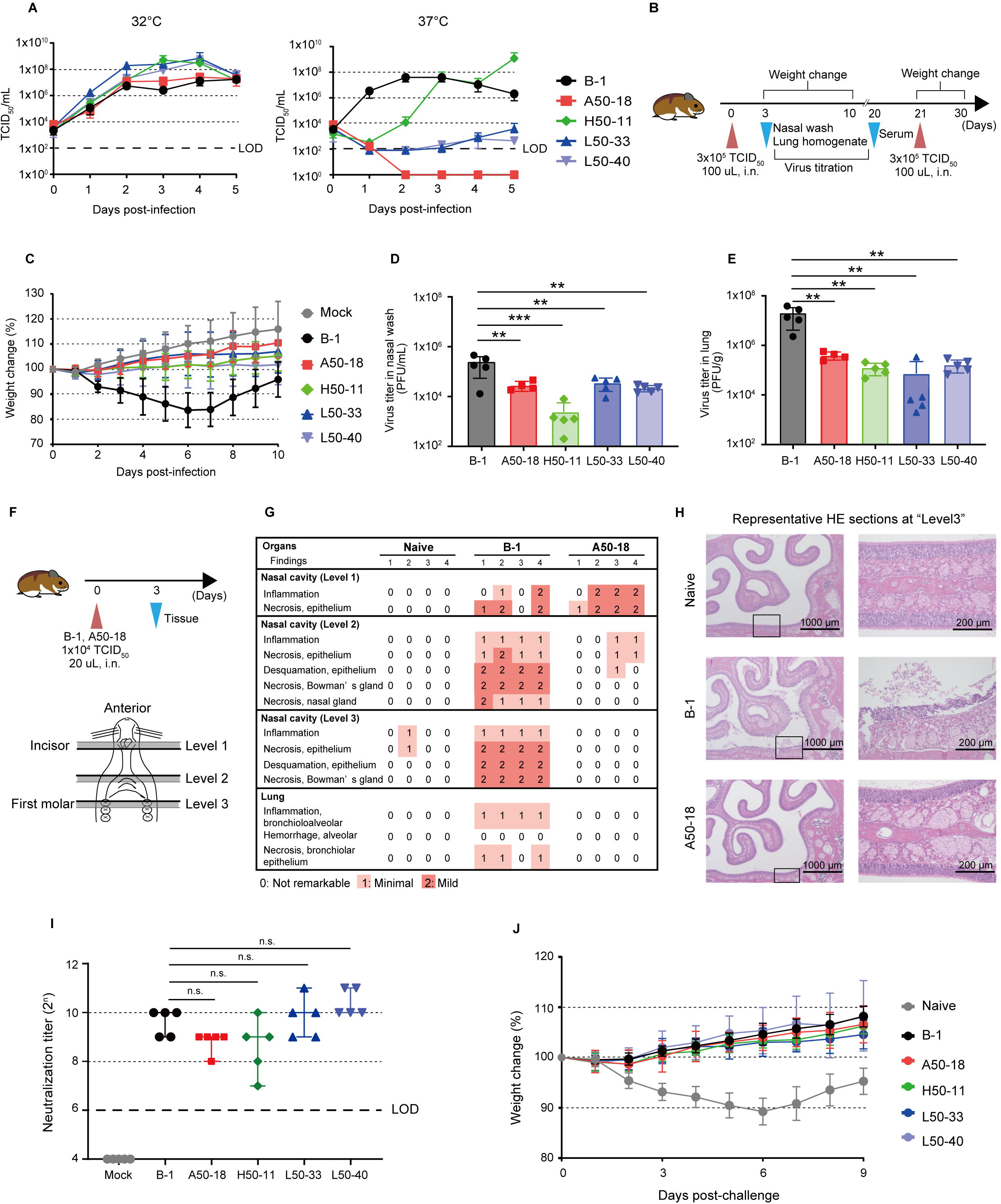
SARS-CoV-2 TS mutants show attenuated phenotype and induce protective immunity. (A) Comparison of the growth kinetics of the four TS mutants with those of the B-1 strain at different temperatures. Vero cells were infected with each strain at multiplicity of infection (MOI) = 0.01. The viral titer of the supernatant was evaluated by TCID_50_ using Vero cells. Symbols represent the average of three independent experiments, and error bars depict the mean SDs. The limit of detection (LOD) is indicated by the bold dashed line. TCID_50_: 50% tissue culture infectious dose, PFU: plaque-forming units. (B) Overview of the evaluation for pathogenicity and immunogenicity of TS mutants (n = 5). (C) Weight changes in the Syrian hamsters infected with the TS mutant or B-1 strains. The average weight change is plotted, and error bars represent the SDs. (D, E) The viral titers of nasal wash specimens (D) and lung homogenates (E) three days post-infection. Infectious viruses were evaluated by plaque formation assays. Bars depict the mean values, and symbols represent individual viral titers. Error bars represent SDs. For statistical analysis, one-way ANOVA was performed (∗∗p < 0.01 and ∗∗∗p < 0.001). (F) Schematic diagram for the evaluation of mucosal tissue damage (n = 4). (G) Damage scores of each section. The percentage of the disrupted area in the entire visual field was classified as 0: not remarkable (< 10%), 1: minimal (10-50%), and 2: mild (50-70%) respectively. (H) Representative hematoxylin and eosin (HE)-stained images of sections at level 3. (I) Neutralization titers of the sera from hamsters infected with each strain and mock-treated hamsters. Median values are plotted, and error bars represent the 95% CIs. For statistical analysis, the Kruskal-Wallis test was performed (n.s.: not significant). (J) Weight change in hamsters after B-1 virus reinfection. The average weight change is plotted, and error bars represent mean SDs.

Next, we evaluated the pathogenicity of TS mutant strains in Syrian hamsters, which are widely used as a model for SARS-CoV-2 infection (Imai et al., 2020; Sia et al., 2020) (Figure 1B). Viral pathogenicity was monitored based on the changes in body weight after viral infection. An approximate decrease in body weight of 15% was observed at six dpi in hamsters infected with the B-1 virus (Figure 1C). In contrast, no body weight loss was observed after infection with any of the tested TS strains. To determine the acute signs that can be observed after infection with the TS strains, B-1- or TS strain-infected Syrian hamsters were euthanized at three dpi, and lung tissue damage was evaluated. The lungs of the B-1-infected hamsters were heavier than those of the mock- or TS-infected hamsters (Figure S1C). Additionally, we observed apparent bleeding and destruction of the alveoli in the lungs of B-1-infected hamsters (Figure S1D). However, we did not observe evident critical tissue damage in the TS-infected hamsters. We also measured the amount of virus remaining in the nasal cavity and lungs at three dpi. The viral titer in the nasal wash specimens of the TS-infected hamsters was significantly lower than that of the B-1-infected hamsters (Figure 1D). The mean viral titers were 2.25 × 10^5^, 2.84 × 10^4^, 2.42 × 10^3^, 3.57 × 10^4^, and 2.14 × 10^4^ PFU/mL in B-1-, A50-18-, H50-11-, L50-33-, and L50-40-infected hamsters, respectively. In addition, the viral titer in the lungs of the TS-infected hamsters was approximately 100 times lower than that in the lungs of the B-1-infected group; the mean titers were 1.87 × 10^7^, 3.98 × 10^5^, 1.26 × 10^5^, 7.25 × 10^4^, and 1.66 × 10^5^ PFU/g in B-1-, A50-18-, H50-11, L50-33, and L50-40-infected hamsters, respectively (Figure 1E). These results suggest that the attenuated phenotype of the TS strains was due to impaired viral replication in the lungs. Moreover, to examine whether infection with the TS strains injures the nasal mucosa, we analyzed transverse cross-sections of the nasal cavity at three dpi (Figure 1F). In the “level 1” section, the anterior sections of the nasal cavity, the mucosal damage observed in A50-18-infected hamsters was similar to that observed in B-1-infected hamsters (Figure 1G). However, in “level 2 and 3” slices (medial and posterior sections, respectively), the tissue damage caused by A50-18 infection was milder than that caused by B-1 (Figure 1H), suggesting that intranasal infection with TS strains hardly injures the inner area of the nasal cavity. To evaluate whether the attenuated TS strains could be used as live vaccines, we measured neutralizing titers in sera collected 20 days after the first infection (Figure 1B). The neutralizing antibody titer was <64, 512-1024, 256-512, 128-1024, 512-2048, and 1024-2048 in the sera of hamsters infected with the mock, B-1, A50-18, H50-11, L50-33, and L50-40 strains, respectively (Figure 1I). Additionally, to assess vaccine efficacy, immunized Syrian hamsters were reinfected with the wild-type B-1 virus 21 days after the first infection (Figure 1B). Primary B-1 strain infection resulted in noticeable body weight loss in naïve hamsters whereas no significant body weight decrease was observed in hamsters pre-infected with B-1 or TS strains (Figure 1J), suggesting the induction of protective immunity. These results indicate that the four novel attenuated TS strains can be used as live vaccines.

### The substitutions nsp3 445F, nsp14 248V plus 416S, and nsp16 67I are crucial for the TS phenotype

We then performed a deep sequencing analysis to identify mutations in the four TS strains (Table 1). The A50-18 strain had six missense mutations in the genes encoding nsp14, S, envelope (E), and nucleocapsid (N) proteins. The H50-11 strain had four missense mutations in the *nsp3*, *nsp16,* and *spike* genes. There were two missense mutations in the *nsp3* gene of the L50-33 strain and three missense mutations in the *nsp3* and *spike* genes of the L50-40 strains. The H50-18, L50-33, and L50-40 strains had a common deletion in *orf7a-orf8* (27549-28251).

**Table 1.** Genetic mutation and amino acid substitution in the TS strains.

| Strain | Genetic mutation | Protein | Amino acid substitution |
| --- | --- | --- | --- |
| A50-18 | G18782T | nsp14 | G248V |
|  | G19285A |  | G416S |
|  | C19550T |  | A504V |
|  | C24198T | Spike | A879V |
|  | T26327C | Envelope | L28P |
|  | C28278T | Nucleocapsid | S2F |
| H50-11<sup>a</sup> | T3930C | nsp3 | V404A |
|  | G8213A |  | D1832N |
|  | G20857A | nsp16 | V67I |
|  | C23778A | Spike | T739K |
| L50-33<sup>a</sup> | C4052T | nsp3 | L445F |
|  | A8094G |  | K1792R |
| L50-40<sup>a</sup> | C4052T | nsp3 | L445F |
|  | A8094G |  | K1792R |
|  | T21723G | Spike | L54W |

<sup>a</sup> H50-11, L50-33, L50-40 strains had a common deletion in *orf7a-orf8* (27549-28251).

To identify TS-related mutations in these TS strains, we sought to obtain revertants of these viruses that can proliferate under high-temperature conditions (37-39°C). Vero cells were infected with each of the TS strains at a high multiplicity of infection (MOI; MOI= 1.0) and incubated at 37, 38, or 39°C. For the A50-18 strain, the number of CPE-positive wells was 19 out of 230 analyzed wells at 37°C and 5 out of 223 wells at 38°C. Sequencing analysis revealed that A50-18 revertants replaced the 248V or 416S substitutions in nsp14 with the wild-type sequence (248G or 416G) but not the other amino acid substitutions (Table 2). Additionally, these revertant viruses proliferated at high temperatures when either of the two substitutions returned to the wild-type amino acid, indicating that the substitutions nsp14 248V and 416G are coordinately involved in the TS phenotype. We compared the predicted crystal structures of A50-18 nsp14 to those of B-1 using Alphafold2 (Jumper et al., 2021). The model suggested that both 248V and 416S are located near the zinc finger2 domain, which has been reported to be important for enzymatic activity (Figure S2A). The 248V substitution prevented the formation of one hydrogen bond, whereas 416S altered the angle of the other. We obtained H50-11 revertants in 2/230 wells at 37°C and in 1/230 wells at 38°C. In these viruses, the nsp16 67I substitution was changed to the wild-type amino acid (67V) (Table 2). The crystal structure models predicted that the V67I substitution did not significantly affect the structure of nsp16 (Figure S2B). L50-33 revertants were obtained in 34/230 wells at 37°C, 39/228 wells at 38°C, and 13/230 wells at 39°C. L50-40 revertants were obtained in 17/230 wells at 37°C, 14/228 wells at 38°C, and 4/216 wells at 39°C (Table 2). The number of wells that produced L50 revertant viruses was higher than that for the A50-18 and H50-11 strains. In the revertants of L50-33 and L50-40, nsp3 445F changed back to the wild-type amino acid (leucine) as expected, or was altered to either cysteine, valine, or isoleucine (Table 2). Taken together, these results suggest that nsp14 248V plus 416S, nsp3 445F, and nsp16 67I are responsible for the TS phenotype of each strain.

**Table 2.** Amino acid substitutions of revertants.

| 37°C |  |  |  |  | 38°C |  |  |  | 39°C |  |  |  |
| --- | --- | --- | --- | --- | --- | --- | --- | --- | --- | --- | --- | --- |
| Parent TS strain | Number of revertants | Amino acid substitution | Codon changes | Number of wells | Number of revertants | Amino acid substitution | Codon changes | Wells | Number of revertants | Amino acid substitution | Codon changes | Number of wells |
| **A50-18** | 19/230 | nsp14 248G | GTG>GGT | 6/19 | 5/223 | nsp14 416G | AGT>GGT | 5/5 | 0/230 | - | - | - |
|  |  | nsp14 416G | AGT>GGT | 13/19 |  |  |  |  |  |  |  |  |
| **H50-11** | 2/230 | nsp16 67V | ATT>GTT | 2/2 | 1/240 | nsp16 67V | ATT>GTT | 1/1 | 0/237 | - | - | - |
| **L50-33** | 34/230 |  | TTT>CTT | 4/34 | 39/228 |  | TTT>CTT | 2/39 | 13/230 |  | TTT>CTT | 2/13 |
|  |  | nsp3 445L | TTT>TTG | 17/34 |  | nsp3 445L | TTT>TTG | 22/39 |  | nsp3 445L | TTT>TTG | 6/13 |
|  |  |  | TTT>TTA | 1/34 |  |  | TTT>TTA | 3/39 |  |  | TTT>TTA | 1/13 |
|  |  | nsp3 445C | TTT>TGT | 8/34 |  | nsp3 445C | TTT>TGT | 7/39 |  | nsp3 445C | TTT>TGT | 2/13 |
|  |  | nsp3 445V | TTT>GTT | 3/34 |  | nsp3 445V | TTT>GTT | 3/39 |  |  |  |  |
|  |  | nsp3 445I | TTT>ATT | 1/34 |  | nsp3 445I | TTT>ATT | 2/39 |  | nsp3 445V | TTT>GTT | 2/13 |
| **L50-40** | 17/230 | nsp3 445L | TTT>CTT | 2/17 | 14/228 | nsp3 445L | TTT>CTT | 2/14 | 4/216 | nsp3 445L | TTT>CTT | 2/4 |
|  |  |  | TTT>TTG | 9/17 |  |  | TTT>TTG | 9/14 |  |  | TTT>TTG | 1/4 |
|  |  | nsp3 445C | TTT>TGT | 6/17 |  | nsp3 445C | TTT>TGT | 3/14 |  | nsp3 445C | TTT>TGT | 1/4 |

### The phenotype and attenuation of the TS strains are attributed to the substitutions nsp14 248V plus 416S, nsp3 445F, and nsp16 67I

To confirm the role of the identified substitutions in the TS phenotype, we constructed recombinant viruses using a circular polymerase extension reaction (Torii et al., 2021) (Figure 2A). Recombinant viruses bearing the 248V and/or the 416S substitutions in the nsp14 protein (r14_248V_, r14_416S_, r14_248V, 416S_, respectively), the 67I substitution in the nsp16 (r16_67I_), the 445F substitution in the nsp3 (r3_445F_), and the deletion in the orf7a-orf8 (rΔORF7a-8) were generated. *In vitro* growth kinetics (Figure 2B) and *in vivo* pathogenicity (Figures 2C and 2D) of the viruses were compared. Under the 32°C culture condition, all recombinant mutants (r3_445F_, r16_67I_, r14_248V_, r14_416S_, r14_248V, 416S_, and rΔORF7a-8) replicated comparably to the recombinant B-1 (rB-1) strain (Figure 2A). The r14_248V_ and r14_416S_ mutants proliferated comparably to rB-1 at 37°C, whereas r14_248V, 416S_ showed relatively slower growth at 37°C, and this difference was more pronounced at 39°C. The replication of r14_248V, 416S_ was impaired at 39°C, whereas that of r14_248V_ was slightly delayed. In contrast, the replication of r14_416S_ was not affected at 39°C. These results suggest that a double amino acid substitution in nsp14 (248V and 416S) is necessary for a stronger TS phenotype, which can be observed at 37°C. The proliferation of the r3_445F_ virus was relatively slower or smaller in scale than that of rB-1 at 37°C. Moreover, at 39°C, the replication of these mutants was impaired, indicating that the 445F substitution in nsp3 is responsible for the TS phenotype. The r16_67I_ virus proliferated comparably to rB-1 at 37°C; however, the peak in viral growth at 39°C was delayed compared to that of rB-1. These results suggest that the 67I substitution in nsp16 is responsible for the TS phenotype. Additionally, rΔORF7a-8, which is a recombinant mutant with a deletion in *orf7a-orf8* common to H50-11, L50-33, and L50-40, showed a relatively slower replication at 39°C. This result suggests that this deletion negligibly accounts for the TS phenotype. The A50-18 strain has an A504V substitution in nsp14 in addition to G248V and G416S. This triple amino acid substitution impaired plaque formation at 37 and 39°C (Figure S3), suggesting that nsp14 504V is also involved in temperature sensitivity, although it is unnecessary.

**Figure 2.**
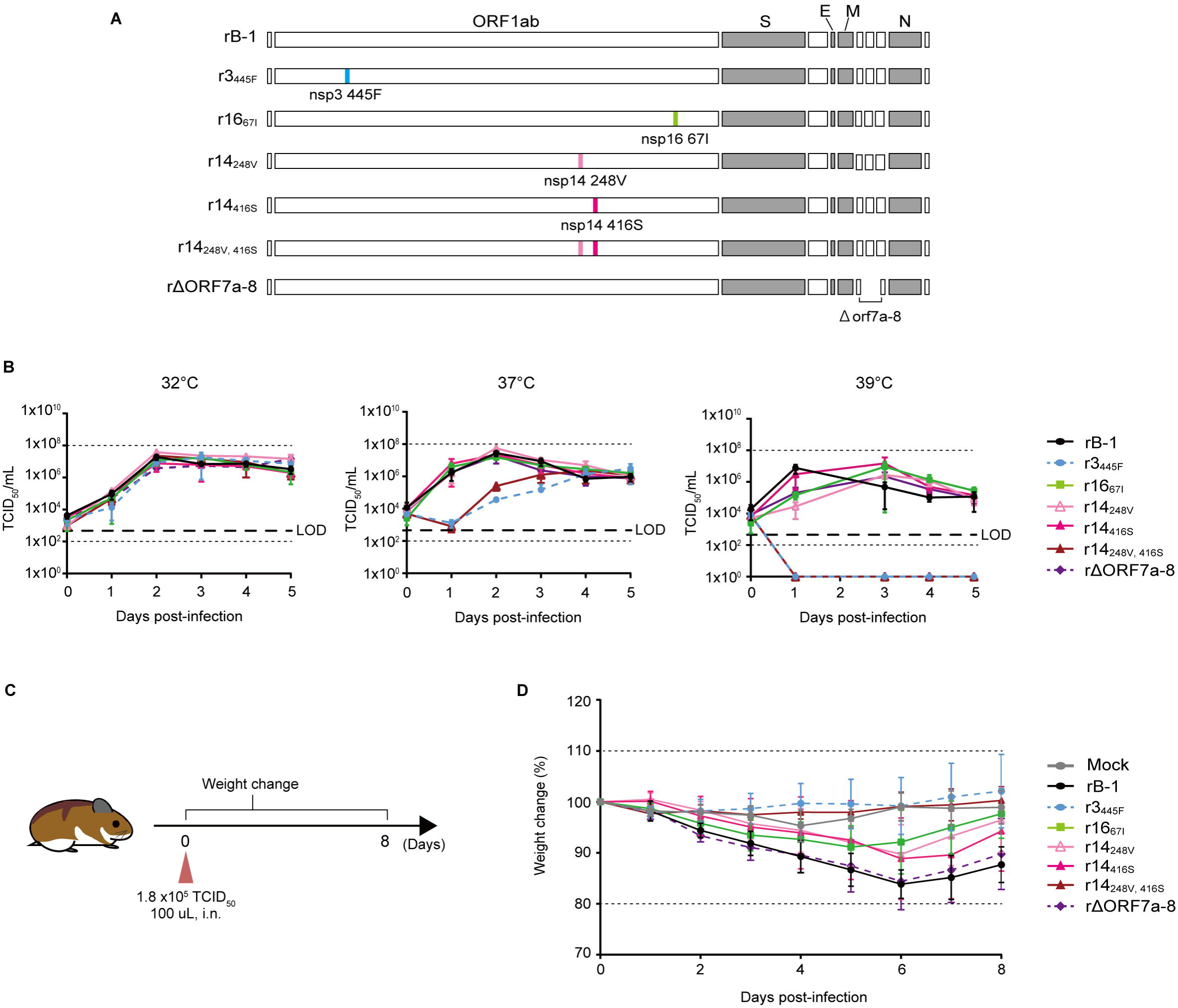
The phenotype and attenuation of the TS strains are attributed to the substitutions nsp14 248V plus 416S, nsp3 445F, and nsp16 67I. (A) The construct of recombinant viruses bearing the substitutions or the deletion found in the TS strains. (B) The growth kinetics of recombinant viruses at different temperatures. Vero cells were infected with each strain at MOI = 0.01. The viral titer of the supernatant was calculated by TCID_50_ using Vero cells. Symbols represent the average of three independent experiments, and error bars mean SDs. The LOD is indicated by the dashed line. (C) Evaluation of the pathogenicity of recombinant viruses *in vivo* (n = 5). (D) Weight change in the Syrian hamsters infected with the recombinant viruses. The average weight changes are represented by symbols, and SDs are represented by error bars.

Next, we assessed whether these recombinant TS mutants were as attenuated *in vivo* as the isolated parental TS strains, using Syrian hamsters (Figure 2C). An approximate body weight decrease of 15% was observed at six dpi in rB-1-infected hamsters (Figure 2D). In contrast, no body weight loss was observed after infection with the r3_445F_ or r14_248V_, _416S_ viruses. The r14_248V_-, r14_416S_-, and r16_67I_- infected groups showed a decrease in body weight of approximately 10%; however, it was significantly less than that observed in the rB-1-infected group. The rΔORF7a-8-infected group lost as much weight as the rB-1-infected group did. Therefore, considering the *in vitro* growth kinetics data, the temperature sensitivity of the mutants was consistent with their level of attenuation.

### The rTS-all vaccine candidate strain is highly attenuated and shows high immunogenicity in Syrian hamsters

As described above, the TS-associated substitutions changed back to the wild-type amino acid when the virus was found to proliferate at high temperatures (Table 2). Moreover, hamsters infected with the L50-33 revertant exhibited a decrease of approximately 21% in body weight at six dpi (Figure S4), suggesting a risk of virulent reversion if TS strains are used as vaccines. To decrease the risk of virulent reversion and to generate a safe vaccine candidate, we combined the TS-associated mutations of *nsp3*, *nsp14*, *nsp16,* and Δ*orf7a-orf8*, whose mechanisms were predicted to be independent of one another (rTS-all). Thus, these mutations may act in synergy or complement each other, rendering a more stable TS phenotype. To assess the reversion risk of rTS-all, we performed the experiment described in Figure 3A. Each TS virus was transferred to 39°C culture conditions, a non-permissive temperature, after an incubation period of one or two days at 32°C. Three days after passaging, we counted the number of CPE-positive wells and checked the genomic sequence of the TS-responsible substitutions of viruses in these wells. Replication of the B-1 strain was observed in all wells (Figure 3B). In contrast, no CPE-positive wells were detected in the rTS-all-inoculated wells, whereas several revertants were detected in the isolated TS strains (A50-18, H50-11, and L50-33). These results suggest that the risk of virulent reversion of rTS-all might be lower than that of the isolated TS strains. Moreover, the *in vitro* growth kinetic assays demonstrated that the replication of rTS-all was impaired under 37°C and 39°C culture conditions (Figure 3C). Even at 32°C, rTS-all started to proliferate later than rB-1, at one dpi. These results suggest that rTS-all is highly sensitive to temperature.

**Figure 3.**
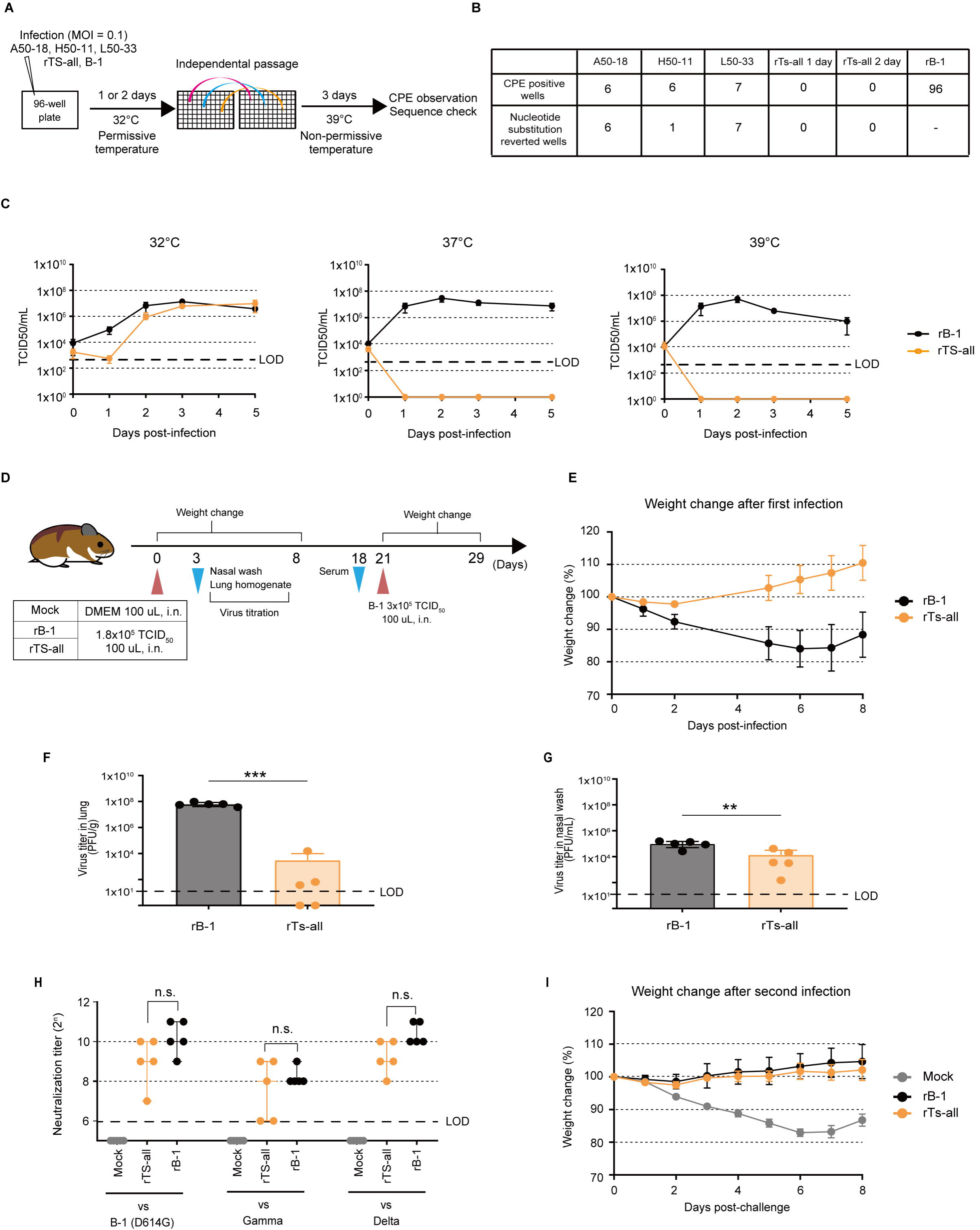
The rTS-all vaccine candidate strain is highly attenuated and shows high immunogenicity in Syrian hamsters. (A) Scheme for the evaluation of the risk of virulent reversion by their ability to replicate at lower temperatures. (B) The number of CPE-positive wells (upper line) and that of those that changed back to the wild-type sequence (lower line). (C) Growth dynamics of the rTS-all and rB-1 strains at different temperatures. Vero cells were infected with each strain at MOI = 0.01. Infectious viruses in the supernatants were evaluated by TCID_50_ in Vero cells. Symbols represent the average of three independent experiments, and error bars represent SDs. The LOD is shown with the bold dashed line. (D) Overview of the investigation of the pathogenicity and immunogenicity of the rTS-all strain (n = 5). (E) Weight change in the recombinant viruses-infected hamsters. The average weight change is plotted, and error bars represent the SDs. (F, G) The viral titers in lung homogenates (F) and nasal wash specimens (G) three days post-infection. Viral titration was performed by plaque formation assays. Bars depict the mean values, and symbols represent individual viral titers. Error bars represent mean SDs. The LOD is indicated by the dashed line. For statistical analysis, one-way ANOVA was performed (∗∗p < 0.01 and ∗∗∗p < 0.001). (H) Neutralization titers in the sera of rB-1 or rTS-all-infected and mock-treated hamsters. Symbols represent individual neutralization titers. Bars show median values and error bars represent 95% CIs. For statistical analysis, the Kruskal-Wallis test was performed (n.s.: not significant). (I) Weight change in Syrian hamsters after reinfection with the B-1 virus. The average weight change is plotted, and error bars represent mean SDs.

To investigate the *in vivo* levels of attenuation and immunogenicity, Syrian hamsters were first inoculated intranasally with 1.8 × 10^5^ TCID_50_/100 µL/dose of rB-1 or rTS-all and reinfected with the wild-type B-1 virus 21 days later (Figure 3D). No body weight loss was observed after primary infection with rTS-all; however, the body weight of the rB-1-infected group decreased by approximately 15%, suggesting that rTS-all was indeed attenuated (Figure 3E). Moreover, the viral titer in the lungs of rTS-all-infected hamsters was remarkably lower than that in the lungs of rB-1-infected hamsters, with mean viral titers of 6.29 × 10^7^ and 3.08 × 10^3^ PFU/g in rB-1- and rTS-all-infected hamsters, respectively (Figure 3F). The viral titer in nasal wash specimens of rTS-all-infected hamsters was also markedly lower than that of rB-1-infected hamsters, with mean viral titers of 1.01 × 10^5^ and 1.39 × 10^4^ PFU/mL in rB-1- and rTS-all-infected hamsters, respectively (Figure 3G). These results indicated that rTS-all is a hyper-attenuated mutant as its replication in the lungs is greatly impaired.

The titer of neutralizing antibodies in the serum of mock-infected hamsters was <64 (Fig. 3H). Infection with rTS-all induced neutralizing antibodies as well as rB-1 did, titer ranges being 128-1024 (vs B-1), 64-512 (vs Gamma strain: hCoV-19/Japan/TY7-501/2021), 256-1024 (vs Delta strain: BK325), respectively. These results suggest that the humoral immune responses induced by infection with the attenuated rTS-all was comparable to that induced by rB-1, even against heterologous virus infection. Importantly, no significant weight decrease was observed after challenging hamsters pre-infected with rB-1 or rTS-all with the B-1 virus (Figure 3I), suggesting that infection with rTS-all induced protective immunity.

### Live-attenuated TS viruses are effective against the variant Omicron

Antigenic drift has enabled Omicron, the latest VOC, to propagate around the world and become dominant in one month (CDC). To evaluate whether rTS-all-infection protects against the Omicron variant, Syrian hamsters were inoculated intranasally with 1.8 × 10^5^ TCID_50_/100 µL/dose of rB-1 or rTS-all. Inoculation with rB-1 and rTS-all induced neutralizing antibodies against the homologous virus B-1 (Figure 4B). Furthermore, titers against the Omicron variant were <32-128 in hamsters infected with rB-1 or rTS-all (Figure 4C). To evaluate whether rTS-all induces heterologous protection, animals were challenged with the TY38-873 Omicron virus (7.2 × 10^4^ TCID_50_/20 µL) 28 days after the initial inoculation (n = 5, Figure 4A). Viral titers in the lungs and nasal specimens, collected three days after Omicron infection, were measured by plaque assays. In the lungs, the mean viral titer in the rB-1- and rTS-all-infected groups was significantly lower than that in the mock-infected group, which had a titer of 6.4 × 10^5^ PFU/g (Figure 4D). The mean viral titer in nasal wash specimens of the rB-1- and rTS-all-infected groups was remarkably lower than that of the mock-infected group (5.2 × 10^3^ PFU/mL; Figure 4E). Additionally, the titers in the lungs and nasal wash of the rB-1- and rTS-all-infected groups were not significantly different, suggesting that rTS-all infection can protect hamsters from an Omicron infection, as well as the wild-type infection, by inducing cross-reactive neutralizing antibodies. Moreover, we compared the neutralizing antibody titers in the nasal wash specimens of the rB-1- and rTS-all-infected groups, collected 28 days after primary infection, using a VSVΔG/Luc-encoding SARS-CoV-2 S-expressing pseudovirus (Tani et al., 2010). The nasal wash specimens of the rTS-all-infected group showed significantly lower luciferase activity than that of the mock group, similar to the rB-1-infected group (Figure 4F). This result suggests that infection of the nasal mucosa with rTS-all can induce neutralizing antibodies to protect against viral entry. Collectively, these data demonstrate that infection with rTS-all induces the production of systemic and mucosal neutralizing antibodies, which can prevent infection with heterologous viruses, similar to the wild-type infection.

**Figure 4.**
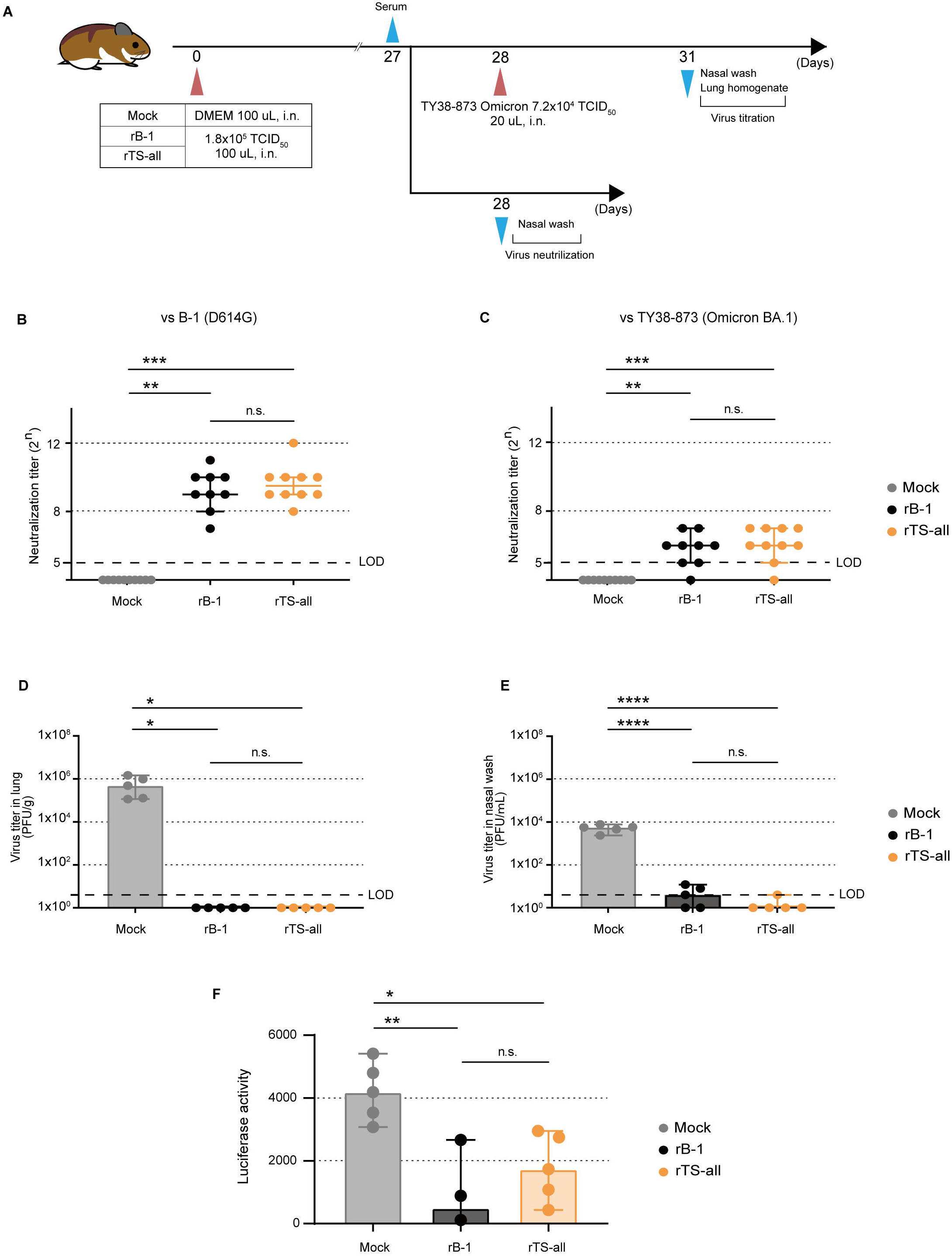
Live-attenuated TS viruses are effective against the variant Omicron. (A) Evaluation of the immunogenicity the rTS-all strain in *vivo*. (B, C) Neutralization titers in the sera of immunized hamsters against the B-1 (B) and BA.1 viruses (C). Individual neutralization titers are plotted with symbols and bars showing the median values (Mock and rTS-all; n = 10, rB-1; n = 9). Error bars represent 95% CIs. The LOD is indicated by the dashed line. Kruskal-Wallis test was used for statistical analysis (n.s.: not significant, ∗∗p < 0.01 and ∗∗∗p < 0.001). (D, E) Viral titers in lung homogenates (D) and nasal wash specimens (E) three days post-infection. Viral titration was performed by plaque formation assays. Bars depict the mean values, and symbols represent individual viral titers (n = 5). Error bars represent mean SDs. The LOD is indicated by the dashed line. For statistical analysis, one-way ANOVA was performed (n.s.: not significant, ∗p < 0.05, and ∗∗∗∗p < 0.0001). (F) Neutralization assay of the nasal wash specimens against pseudotyped VSV with D614G pre-alpha type spike. The symbols represent individual data and error bars represent SDs (Mock and rTS-all; n = 5, rB-1; n = 3). The data were statistically analyzed using one-way ANOVA (n.s.: not significant, ∗p < 0.05 and ∗∗p < 0.01).

### A strategy based on TS-associated mutations and adjusted with tailor-made S proteins to target different variants

The identified TS substitutions were located in the viral nsps and did not affect the antigenicity of the S proteins. Thus, we hypothesized that we could design a vaccine platform against VOCs by exchanging the S protein of rTS-all with that of the other strains. To confirm this hypothesis, we constructed a recombinant virus (rTS-all-Omicron) containing the coding sequence of the Omicron S protein within the TS-all backbone (Figure 5A). As expected, rTS-all-Omicron exhibited a TS phenotype *in vitro* (Figure 5B). Furthermore, to evaluate its efficacy as a vaccine against Omicron, we measured the titer of neutralizing antibodies in the serum of hamsters 14 days after infection (Figure 5 C). Consistent with our findings in Figure 4C, the titer against Omicron in the rTS-all-infected group was low (<32 at 14 dpi). However, infection with the rTS-all-Omicron significantly increased the titer of neutralizing antibodies against Omicron (128-256) but not that against the B-1 strain (<32), which has the D614G S protein (Figure 5D). These results confirmed that vaccination with rTS-all-Omicron is effective at protecting against Omicron, suggesting the possibility that rTS-all constitutes a powerful platform for the development of a COVID-19 live-attenuated vaccine by replacing the spike protein with that of newly emerging variants.

**Figure 5.**
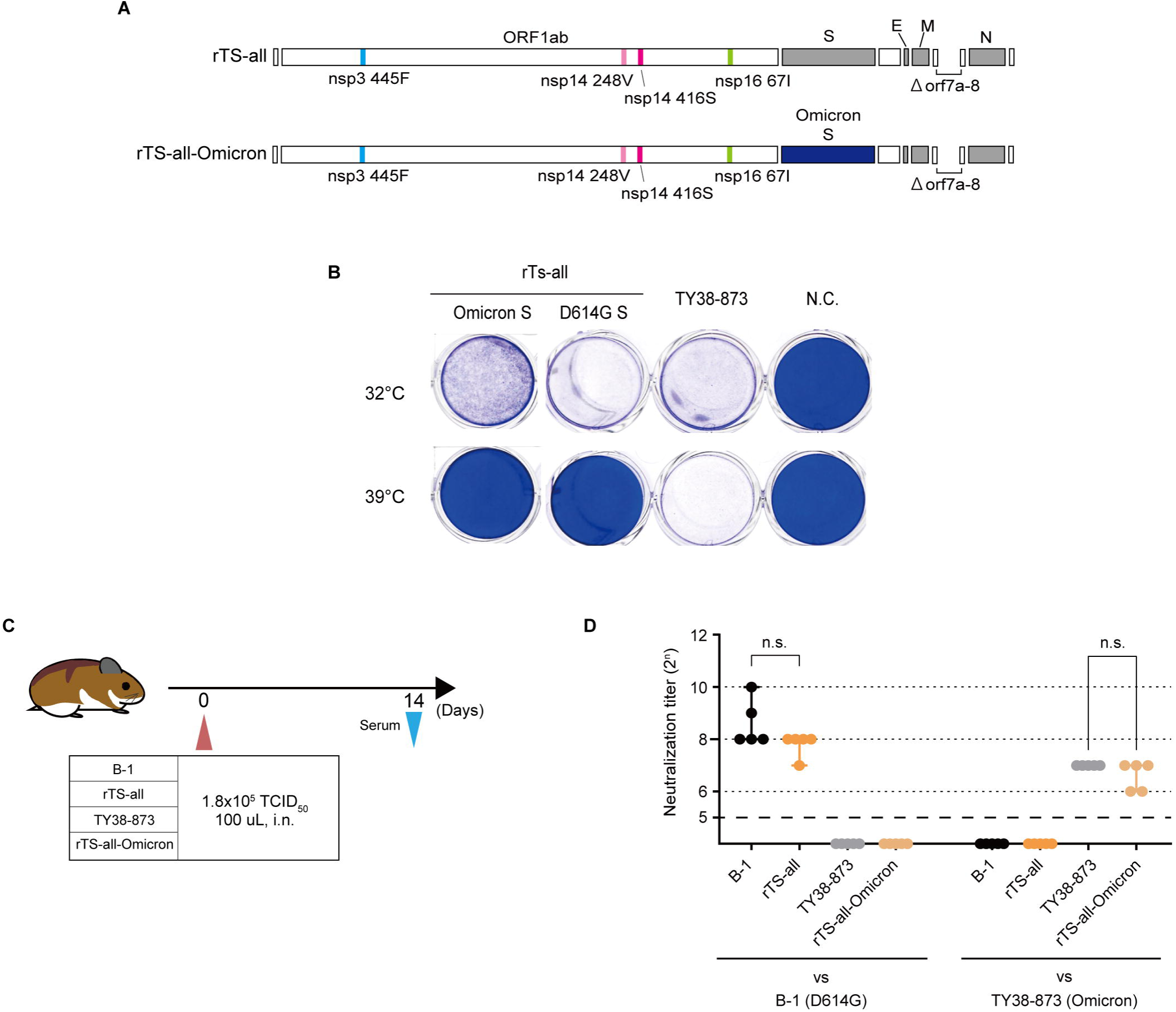
A strategy based on TS-associated mutations and adjusted with tailor-made S proteins to target different variants. (A) The construct of recombinant viruses which contained the Omicron spike with all the substitutions or deletions involved in temperature sensitivity. (B) Temperature sensitivity of rTS-all-Omicron *in vitro*. Vero cells were infected with the recombinant strains and the Omicron variant and incubated under the indicated temperature conditions. CPE were observed three dpi by formalin fixation and crystal violet staining. (C) Evaluation of the immunogenicity of the rTS-all-Omicron strain *in vivo* (n = 5). (D) Neutralization titer of the sera from the hamsters infected with each strain. The individual neutralization titers are indicated with the corresponding symbols. Bars represent median values and error bars mean 95% CIs. For statistical analysis, the Kruskal-Wallis test was performed (n.s.: not significant).

## DISCUSSION

Live-attenuated viruses constitute a highly effective vaccine modality that has been used to treat various infectious diseases. They have several advantages including the induction of an effective humoral immune response and a long-lasting protective cellular immunity without the need for multiple doses. However, there are concerns regarding virulence reversion, such as that reported for oral poliovirus vaccines (Kew et al., 2005). In this study, we identified novel TS-associated mutations in several SARS-CoV-2 nsps that resulted in TS phenotypes of varying degrees of *in vitro* and *in vivo* attenuation in the Syrian hamster model (Figures 1 and S1B-D). To develop a safe and effective live-attenuated vaccine, we hypothesized that combining all these mutations may lead to a virus that is less likely to revert to the more pathogenic wild-type phenotype. Thus, we generated a recombinant virus, rTS-all, and confirmed that it retained TS characteristics *in vitro* and *in vivo*. More importantly, the generation of revertants proved to be less likely in comparison, strengthening the potential use of rTS-all as a vaccine against COVID-19 (Figure 3).

One of the identified mutations resulted in a V67I substitution within the viral nsp16 protein, which is a 2-O-methyltransferase involved in viral genome replication (Chen et al., 2011; Decroly et al., 2011; Decroly et al., 2008; Krafcikova et al., 2020). Structural analyses (AlphaFold2; Figure S2B) (Jumper et al., 2021) of this protein predicted no drastic differences compared with the wild-type version, consistent with the weak phenotype of the strain (H50-11). Another substitution was found in the MACS domain of nsp3 (L445F), accounting for the modest TS phenotypes of the L50-33 and L50-40 strains. In mouse hepatitis virus (MHV), mutations in the nsp3 MAC domain can cause temperature sensitivity, likely by enhancing proteasome-mediated degradation (Deng et al., 2019). Therefore, we speculated that the mechanisms underlying the TS phenotype in our isolated strains might be similar. Finally, several substitutions were identified in nsp14 (G248V, G416S, and A504V), rendering the resulting A50-18 strain more sensitive to higher temperatures. The predicted structure model suggested that both 248V and 416S substitutions affected hydrogen bonds (Figure S2A). The nsp14 protein has exo-ribonuclease and N7 methyltransferase domains and plays an important role in viral genome replication by forming a complex with other nsps (Chen et al., 2009; Minskaia et al., 2006; Ogando et al., 2020). Therefore, at high temperatures, these structural changes may affect the molecular interactions within nsp14 itself and with other nsps, resulting in a decline in viral genome replication.

The strain, rTS-all, which combines all the above mutations, exhibited an attenuated phenotype while maintaining robust immunogenicity *in vivo* (Figure 3). Infection with rTS-all induced serum neutralizing antibodies against not only the homologous virus but also heterologous viruses, that were comparable to those induced by rB-1 infection (Figures 3H, 4 B, and 4C). These results are most likely due to a more complex immunization process. Despite attenuation, the infecting virus stimulates several pathways of the immune system throughout its cycle within the infected cells (Shah et al., 2020), which might induce a highly effective immune response. A previous study has suggested that infection-induced primary memory B cells undergo more affinity maturation than vaccine-induced memory B cells do (Pape et al., 2021). Therefore, rTS-all infection may also induce mature B cells.

A previous study has reported that mucosal IgA against SARS-CoV-2 is present in convalescent patients and contributes to virus neutralization (Sterlin et al., 2021). However, it is not induced after vaccination with mRNA through the intramuscular route (Piano Mortari et al., 2021). An animal model study has reported that vaccination-induced systemic neutralizing antibodies failed to protect nasal tissue against SARS-CoV-2 infection (Zhou et al., 2021). Consistent with these studies, in the present study, we observed that intranasal administration of live-attenuated rTS-all induced neutralizing antibodies not only in the serum but also in the nasal mucosa, similar to the wild-type infection (Figures 4B, 4C, and 4F). Moreover, rTS-all-infected animals were protected from not only the B-1 strain but also the Omicron variant (Figures 3I, 4D, and 4E). We speculate that mucosal neutralizing antibodies induced by rTS-all intranasal administration might play an important role in protection against SARS-CoV-2 in peripheral areas.

A serious concern related to intranasal administration of live viruses is olfactory dysfunction, which has been reported after infection with SARS-CoV-2 (Giacomelli et al., 2020), probably as a consequence of nasal tissue damage (Khan et al., 2021; Urata et al., 2021). In this study, we confirmed that infection with A50-18 resulted in dramatically lower damage to the nasal tissue compared to infection with the wild-type strain. Therefore, our data suggest that the intranasal administration of rTS-all might constitute a safe option for the development of an effective vaccine strategy for protection against SARS-CoV-2 infections.

Previous studies have suggested that SARS-CoV-2 induces a broad, robust, and specific T-cell response in convalescent individuals (Grifoni et al., 2020; Sekine et al., 2020). In our study, rTS-all exhibited a lower proliferation rate than the wild-type virus in the lungs, as well as less pathogenicity (Figure 3F). However, it replicated in the nasal cavity, which is a low-temperature region (Figure 3G). Although we did not assess cellular immunity in this study, infection with the rTS-all strain might elicit a T-cell response similar to that of the wild-type strain.

Our study also offers a strong vaccine platform applicable to future variants. As nsp3, nsp14, and nsp16 do not affect the antigenicity of the S protein, alternative TS strains can be generated by changing the S coding sequence. Here, we confirmed that rTS-all-Omicron exhibited immunogenicity against the Omicron variant and conserved the TS phenotype characteristics of the original rTS-all strain. Notably, we were able to generate the rTS-all-Omicron virus two weeks after the Omicron sequence was made available. Overall, our data suggest that this platform could be a promising candidate for the development of a live-attenuated vaccine that can be rapidly produced in response to new emerging variants or another SARS-CoV pandemic in the future.

### Limitations of the study

Detailed characterization of the cellular and humoral immune responses was difficult using this model, due to the lack of hamster-specific reagents. Using a genetically modified mouse model could help circumvent this problem. Furthermore, it is unclear whether virulent reversion occurs during *in vivo* transmission, thus additional studies are required. In addition, olfactory functional analysis could help better determine its correlation with the nasal tissue injury, an important point for the development of intranasal attenuated vaccines.

## ACKNOWLEDGMENTS

We appreciate the assistance from Mitsuyo Kosaka, Mai Matsumoto, and Manae Morishima (BIKEN). The authors acknowledge the NGS core facility of the Genome Information Research Center at the Research Institute for Microbial Diseases of Osaka University for their support with RNA-seq. This work was supported by The Research Foundation for Microbial Diseases of Osaka University (BIKEN) and Japan Agency for Medical Research and Development (AMED) under Grant Number JP20pc0101047.

## AUTHOR CONTRIBUTIONS

H.E. conceived and designed the study. A.Y. and S.O. conducted the majority of the experiments. S.T., C.O., and Y.M. constructed the CPER fragments for the construction of recombinant viruses. S.K. performed the pseudovirus neutralization assays. H.S., S.U., and W.K. provided scientific insight. A.Y., S.O., and P.M. wrote the original manuscript and H.E. revised it. S.T., K.Y., and H.E. supervised the project.

## RESOURCE AVAILABILITY

### Lead contact

Further requests for resources and reagents should be directed to and will be fulfilled by the lead contact, Hirotaka Ebina.

### Materials availability

The principal authors of this paper are employees of the Research Foundation for Microbial Diseases of Osaka University (hereafter, the Foundation). Therefore, the distribution of the TS strains and their composite strains obtained in this study requires the establishment of an MTA with the Foundation.

### Data and code availability

The data in this report are available from the lead contact upon request. The viral genome sequences of the TS mutants isolated in this study have been deposited at the National Center for Biotechnology Information (Accession numbers 5 for B-1, A50-18, H50-11, L50-33 and L50-40 are LC603286, LV603287, LC603288, LC603289 and LC603290, respectively). All other data needed to evaluate the conclusions in the paper are present in the paper or the Supplemental information. Additional information for reanalysis is shared by the lead contact upon request.

## EXPERIMENTAL MODEL AND SUBJECT DETAILS

### Ethical statement

All animal experimental protocols, including anesthesia conditions, endpoints for infection, and euthanasia methods, were reviewed and approved by the Osaka University Animal Experiment Committee.

### Cell lines

Vero cells (African green monkey kidney cells: ATCC, CCL-81) and 293T cells (human embryonic kidney cells: ATCC, CRL-3216) were maintained at 37°C in Dulbecco’s modified Eagle’s Medium (DMEM, Sigma-Aldrich, Cat# D6429) supplemented with 10% heat-inactivated fetal bovine serum (FBS, Sigma-Aldrich), penicillin (100 U/mL), and streptomycin (0.1 mg/mL) (PS, Gibco, Cat# 1154887). VeroE6-TMPRSS2 cells (TMPRSS2-expressing Vero cells: JCRB, JCRB1819) were maintained at 37°C in DMEM supplemented with 10% heat-inactivated FBS, PS, and 1 mg/mL G418 Sulfate (Gibco, Cat# 10131027). BHK cells (hamster kidney cells: JCRB, JCRB9020) were maintained at 37°C in Minimum Essential Medium Eagle (MEM, Sigma-Aldrich, Cat# M4655) supplemented with 10% heat-inactivated FBS and PS. BHK cells stably expressing human ACE2 were generated by piggyBac plasmid transfection and puromycin-based selection and maintained with 3 µg/mL puromycin (Gibco, Cat# A1113803).

### Viruses

SARS-CoV-2 was isolated from a SARS-CoV-2-positive clinical specimen in Osaka City, Japan. An aliquot of 500 µL of the specimen was diluted with 500 µL of Dulbecco’s phosphate-buffered saline (D-PBS, Nacalai Tesque, Cat# 14249) and filtered through a 0.22 µm filter. One hundred microliters of this filtrate were used to inoculate the Vero cells. Cytopathic effects (CPE) were observed three days post-infection (dpi). The supernatant was collected and stored at −80°C as the “SARS-CoV-2 clinical isolate strain B-1”. Whole-genome sequencing of B-1 was performed using next-generation sequencing (NGS) analysis, as described below. The sequencing results showed that the B-1 strain had a D614G substitution in the S protein. The SARS-CoV-2 delta variant (BK325) was provided by the Research Foundation for Microbial Diseases at Osaka University. The gamma (hCoV-19/Japan/TY7-501/2021, GISAID ID: EPI_ISL_833366) and Omicron variants (TY38-873, GISAID ID: EPI_ISL_7418017) were obtained from the National Institute of Infectious Diseases of Japan (NIID).

A pseudotyped vesicular stomatitis virus (VSV) with the SARS-CoV-2 S protein (pre-alpha type: D614G) was constructed for neutralization assays. The pCAGGS plasmid encoding the codon-optimized SARS-CoV-2 *spike* gene lacking the C-terminal 19 amino acids was transfected into 293T cells using polyethylenimine (Polysciences, Cat# 24765). Twenty-four hours post-transfection, VSVΔG/Luc-complemented VSV G pseudovirus, where the VSV G gene was replaced by the luciferase gene but expressed VSV G on the surface of the virion (VSVΔG/Luc complemented with VSV G), was used to infect cells at multiplicity of infection (MOI) = 0.1 (Tani *et al*., 2010). The excess unattached virus was eliminated by removing the supernatant two hours post-infection and replacing it with a fresh medium. After 72 h of incubation at 37°C, the cell culture supernatants containing VSVΔG/Luc-encoding SARS-CoV-2 S-expressing pseudovirus were collected and filtered through a 0.45 µm pore size filter.

### Evaluation of pathogenicity in hamsters

Syrian hamsters (Slc:Syrian) were purchased from Japan SLC, Inc. Five-week-old male hamsters were anesthetized with 2.0-3.0% isoflurane (FUJIFILM Wako, Cat# 1349003) inhalation and 0.3 mg/kg medetomidine (Nippon Zenyaku Kogyo) + 4 mg/kg midazolam (Maruishi Pharmaceutica) + 5 mg/kg butorphanol (Meiji Seika Pharma) intraperitoneal injection. One hundred microliters of DMEM containing the respective SARS-CoV-2 strains were delivered dropwise to the nostrils, and body weight was measured daily. For the assessment of acute symptoms, lung and nasal wash specimens were collected with 1 mL of D-PBS at three dpi. The lungs were divided into left and four right lungs. The left lung was fixed with 10% formalin and serially sectioned. One section was stained with hematoxylin (Sakura Finetek Japan, Cat# 9130-4P) and eosin (FUJIFILM Wako, Cat# 051-06515). The right lung was homogenized with Biomasher II (Nippi, Cat# 320 103) and suspended in 1 mL of DMEM. After centrifugation at 100 *g* for 5 min, the supernatant was collected as lung homogenate. The viral titers of these samples were evaluated using a plaque formation assay. To evaluate the tissue damage in the nasal cavity caused by infection with the TS mutants, hamsters were infected with SARS-CoV-2 B-1 or A50-18 strains contained in 20 µL of DMEM via the intranasal route to limit the administration to the upper respiratory tract. After euthanasia, the heads were fixed in 10% formalin and sectioned. Each section was stained with hematoxylin and eosin (HE).

### Evaluation of antigenicity in hamsters

To analyze antigenicity, we performed a re-challenge assay and evaluated the presence of neutralizing antibodies in the serum and nasal wash specimens. Hamsters who recovered from the SARS-CoV-2 primary infection were re-challenged with DMEM containing the respective SARS-CoV-2 strains three weeks after the first infection. Body weight changes were recorded for an additional 8-10 d. For the neutralization assay, blood and nasal wash specimens were collected from the recovered hamsters. The blood specimens were centrifuged at 800 *g* for 10 min, and serum was collected for analysis. Nasal wash specimens were filter-sterilized prior to the assay.

## METHOD DETAILS

### Plasmids

The *hACE2* gene was cloned into the piggyBac plasmid (Systembiosciences, PB514B-2). Briefly, total RNA was extracted from 293T cells and reverse-transcribed using the SuperScript™ III First-Strand Synthesis System for RT-PCR (Invitrogen, Cat# 11904018). Then, hACE2 complementary DNA (cDNA) was amplified using the KOD One PCR Master Mix -Blue- (TOYOBO, Cat# KMM-201). The obtained fragment was digested with XbaI (NEB, Cat# R0145) and NotI (NEB, Cat# R0189) and ligated with PB514B-2, which was also digested with XbaI and NotI, using the TAKARA Ligation kit ver.2.1 (TAKARA Bio, Cat# 6022).

### Isolation of TS mutants

To obtain TS mutants, a previously reported protocol for the induction of mutations in MHV was followed (Deng et al., 2019). Briefly, the clinical isolate strain B-1 was used to infect confluent Vero cells in six-well plates for 1 h at 37°C. A fresh medium containing 5-fluorouracil (100 µg/mL, FUJIFILM Wako, Cat# 068-01401) was added. After a one-day incubation period at 32°C, the supernatants were collected and stocked as “mutated virus”. These viruses were passaged three times in Vero cells at 32°C, and viral clones were obtained by plaque isolation. The collected plaques were suspended in 100 µL DMEM and an aliquot of 2-50 µL of this suspension was used to infect Vero cells at 32°C. To confirm temperature sensitivity, we observed the development of CPE at 37 or 32°C.

### Construction of recombinant viruses using circular polymerase extension reaction (CPER)

Viruses bearing TS mutations were constructed using circular polymerase extension reaction (CPER) (Torii et al., 2021), with minor modifications. We cloned the fragmented B-1 viral genome into plasmids. TS mutations of interest were introduced into these plasmids using inverse PCR. We then assembled these fragments by CPER using PrimeSTAR GXL DNA polymerase (Takara Bio, Cat# R050A). The assembled cDNA was transfected into BHK-hACE2 cells using Lipofectamine LTX Reagent with PLUS™ Reagent (Invitrogen, Cat# 15338100). Seven days after transfection, supernatants were collected. The supernatant was used to inoculate VeroE6-TMPRSS2 and incubated for four days. After observing CPE, the supernatant was collected and used to inoculate Vero cells to harvest the desired SARS-CoV2 viruses. To obtain TS-recombinant viruses, we performed a construction experiment at 32-34°C.

### Titration assay

Viral titration was determined using the 50% tissue culture infectious dose (TCID_50_) or plaque formation units (PFU). Briefly, samples were serially diluted in DMEM supplemented with 2% FBS and antibiotics. Fifty microliters of diluted samples were used to infect confluent Vero cells in 96-well plates and incubated at 37°C (wild-type strain) or 32°C (TS strain) for six days. Infected cells were fixed with 10% formalin and stained with crystal violet. After staining, the TCID_50_ was calculated using the Behrens-Karber method. For the plaque formation assay, diluted samples were used to infect confluent Vero cells in 6-well plates for 1 h at 32 or 37°C. Cells were then washed with D-PBS. After washing, a fresh medium containing 1% SeaPlaque Agarose (Lonza, Cat# 50100) was layered and incubated at 32 or 37°C until plaque formation. Cells were fixed in 10% formalin and stained with crystal violet. Visible plaques were counted to calculate the PFU.

### NGS analysis

Vero cells were infected with wild-type or TS SARS-CoV-2 and incubated at 37 or 32°C, respectively. After three days, supernatants were collected, and RNA was extracted using a QIAamp Viral RNA Mini Kit (Qiagen, Cat# 52904) according to the manufacturer’s protocol. Viral RNA was processed and analyzed at the Genome Information Research Center of Osaka University using NovaSeq6000 (Illumina). Wuhan strain (NC045512) was used as the reference sequence.

### Determination of viral growth dynamics

One million Vero cells per well were cultured in six-well plates and incubated at 37°C overnight. SARS-CoV-2 strain suspensions (1 × 10^4^ TCID_50_) were used to inoculate the cells (MOI = 0.01), which were then incubated at 32, 34, or 37°C for five days. The supernatants were collected and stored at −80°C daily. Viral titration of the supernatants was performed using TCID_50_/mL, as described above. Each experiment was performed in triplicate.

### Neutralization assay

Serum samples were inactivated by incubation at 56°C for 30 min and serially diluted with DMEM supplemented with 2% FBS. Serially diluted serum was mixed with 100 TCID_50_ of SARS-CoV-2 B-1, Delta, Gamma, or Omicron strains and incubated at 37°C for 1 h. After incubation, the samples were used to inoculate confluent Vero cells in 96-well plates and were incubated at 37°C for seven days. Cells were fixed in formalin and stained with crystal violet. Neutralizing titers were determined as the inverse of the maximum dilution that prevented viral proliferation.

For the neutralization assay of nasal wash specimens, we performed an assay using the pseudotyped VSV virus. Briefly, the VSVΔG/Luc-encoding SARS-CoV-2 S-expressing pseudovirus was mixed and co-incubated with the same volume of nasal wash and incubated for one hour. Mixtures were added to Vero cells and the cells were lysed using lysis buffer (Promega, Cat# E153A) after incubation for 24 h. Luciferase activity was measured using a ONE-Glo™ EX Luciferase Assay System (Promega, Cat# E8130).

### Generation and characterization of revertants

To obtain revertants, TS strains were used to infect confluent Vero cells at 37, 38, and 39°C (MOI = 1). Viruses from the CPE-positive wells were further propagated in Vero cells at 37 or 38°C to obtain the corresponding viral stocks. To identify any mutations around the nucleotide of interest in the revertants, viral genomic RNA was extracted using TriReagent (Molecular Research Center, Cat# TR118) or a QIAamp viral RNA mini kit (Qiagen, Cat# 52904), according to the manufacturer’s protocols, and subsequently subjected to cDNA synthesis using the SuperScript™ III First-Strand Synthesis System for RT-PCR (Invitrogen, Cat# 11904018). The genomic regions surrounding each mutation of interest in Table 1 were amplified by PCR using specific primers. The PCR products were sequenced and compared with those of the corresponding parental TS strains.

### Estimating the “reversion-to-virulence” risk of each mutant

To estimate the risk of each TS strain reverting to the virulent phenotype, each TS virus was used to infect confluent Vero cells (5 × 10^3^ TCID_50_/well) in 96-well plates, which were maintained for one or two days at 32°C. Subsequently, they were passaged independently in new confluent Vero cells in 96-well plates and maintained for additional three days at 39°C. After incubation, the number of CPE-positive wells was counted. Eventually, we confirmed the viral sequence, as described above.

## QUANTIFICATION AND STATISTICAL ANALYSIS

Each data point is expressed as the mean ± SD or median ± 95% CI. To analyze the viral titer in the nasal wash or lung samples, statistical analyses were performed using one-way ANOVA. For the analysis of the neutralization titer, the Kruskal-Wallis test was used to calculate statistical significance. All analyses were performed using the GraphPad Prism software (n.s.: not significant, ∗p < 0.05, ∗∗p < 0.01, ∗∗∗p < 0.001, and ∗∗∗∗p < 0.0001).

## SUPPLEMENTAL INFORMATION

**Supplementary Figure 1.**
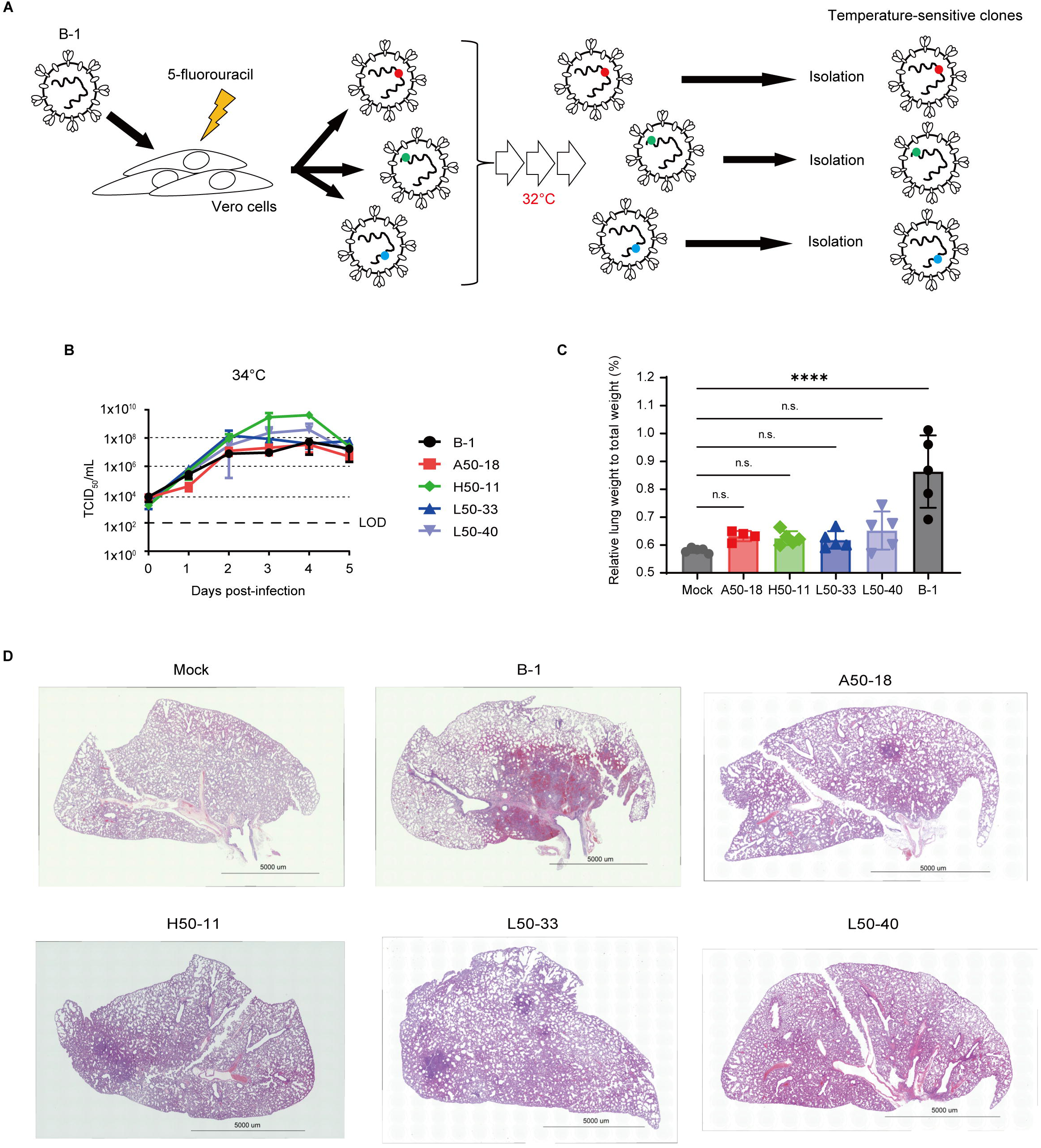
Characterization of the TS strains (Relating to Figure 1) (A) Scheme of TS mutant isolation. (B) The growth kinetics of four TS mutants and B-1 strain at 34°C. Vero cells were infected with each strain at MOI = 0.01. The viral titer of the supernatant was evaluated by TCID_50_ using Vero cells. Symbols represent the average of three independent experiments, and error bars represent SDs. The limit of detection (LOD) is indicated by the bold dashed line. (C) The relative lung weights of hamsters infected with each strain three days post-infection were compared to those of mock-treated hamsters. The average is shown with bars and individual data is represented with symbols. Error bars mean SDs. For statistical analysis, one-way ANOVA was performed (n.s.: not significant and ***p < 0.0001). (D) Representative hematoxylin and eosin (HE)-stained images of lung sections from each TS strain-infected hamster. Scale bar, 5000 µm.

**Supplementary Figure 2.**
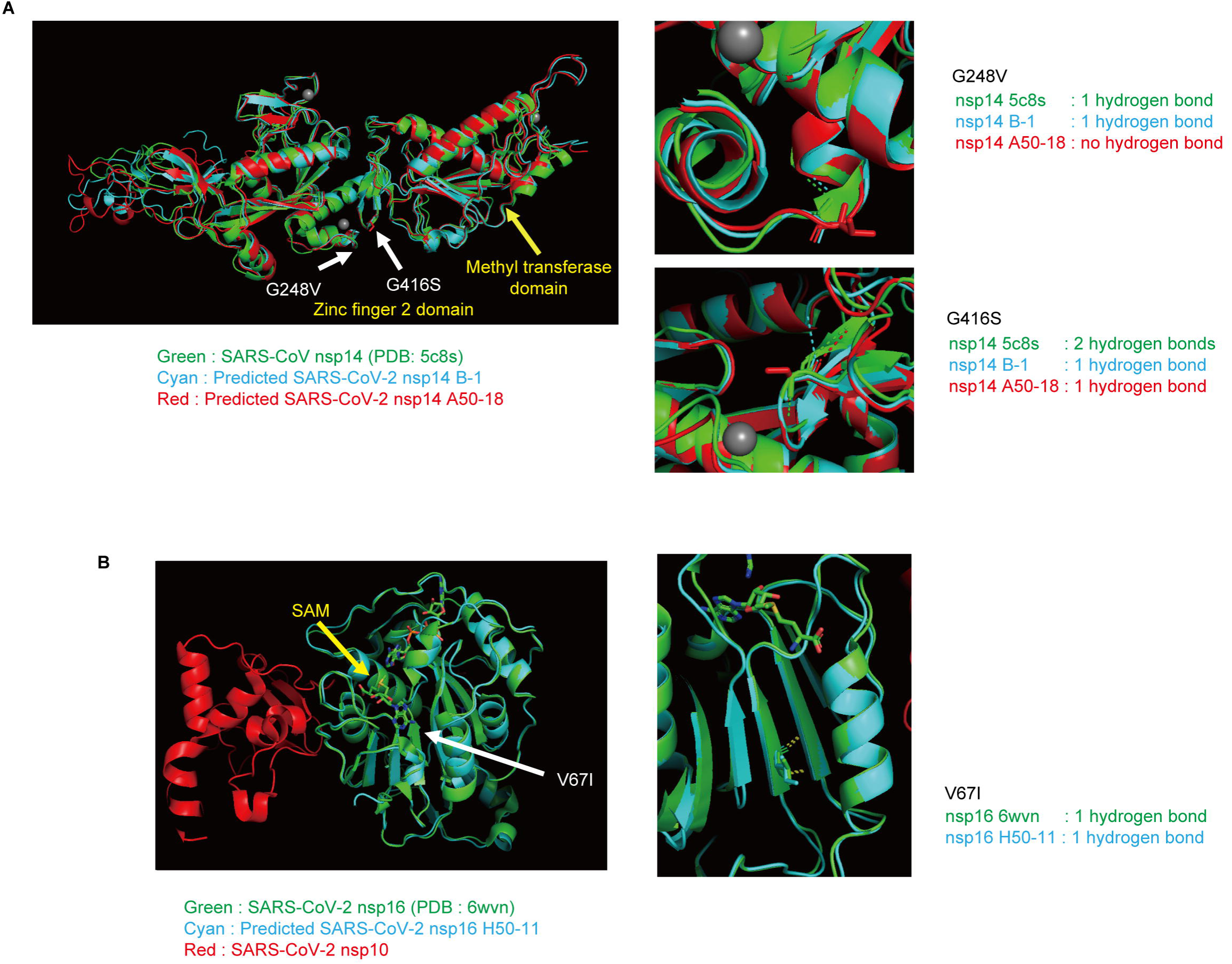
Prediction of the structures of nsp14 and nsp16 with the TS-associated substitutions. (A) The nsp14 structures of the parental (B-1: cyan) and TS strains (A50-18: red) were estimated using Alphafold2 and compared to that of SARS-CoV (PDB: 5c8s, green). Domains are indicated by yellow arrows and TS-associated substitutions are indicated by white allows. (B) The structure of nsp16 of H50-11 (cyan) was predicted using Alphafold2 and compared to that of SARS-CoV-2 (PDB: 6wvn, green). The S-adenosylmethionine (SAM) entry site is shown by the yellow arrow and V67I substitution is indicated by the white arrow.

**Supplementary Figure 3.**
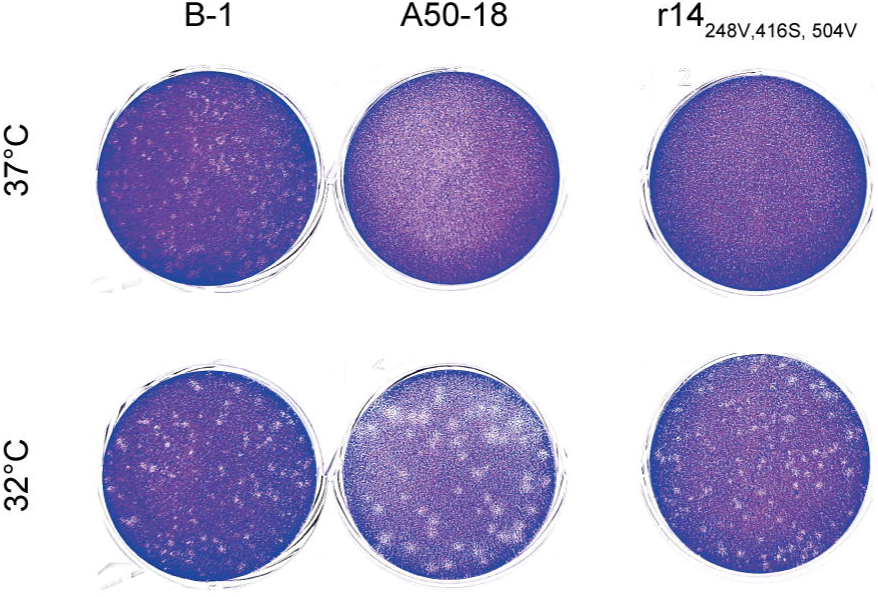
The temperature-sensitive phenotype of r14_248V,416S,504V_ (Relating to Figure 2) Representative images of plaques generated by each strain after a four-day incubation period at the indicated temperatures.

**Supplementary Figure 4.**
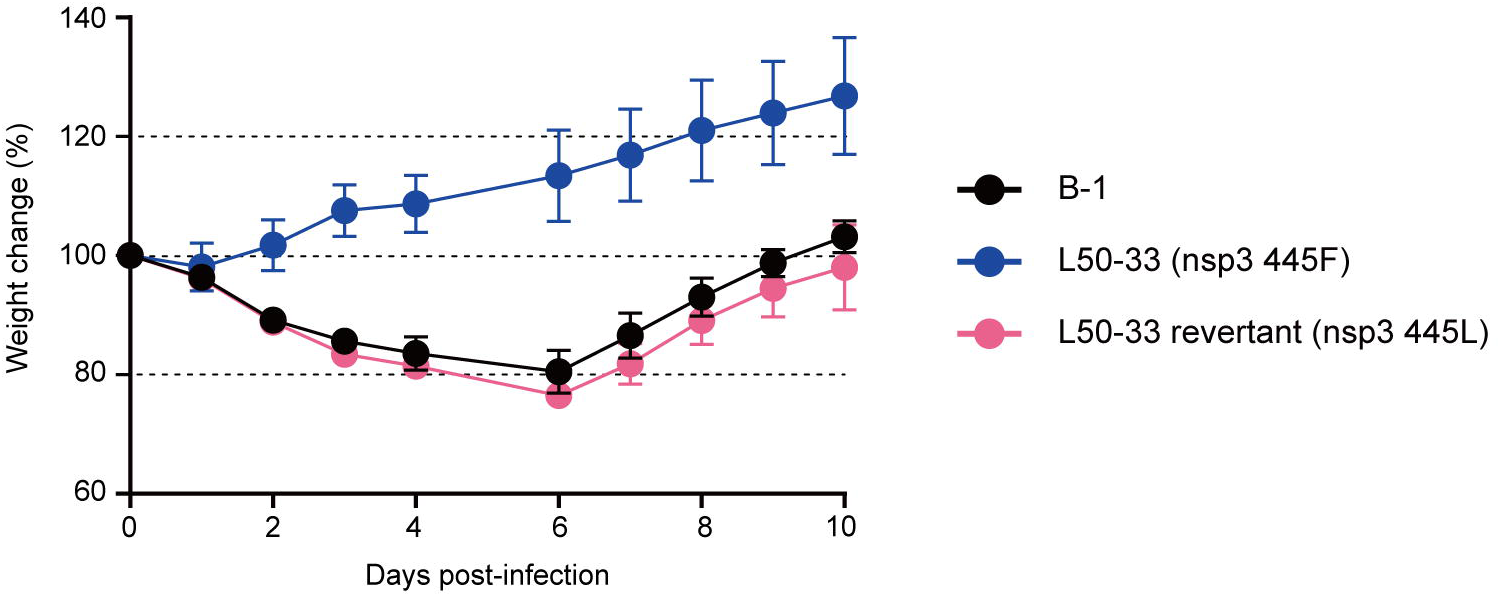
The *in vivo* pathogenicity of the revertant that emerged from L50-33 (Relating to Figure 3) B-1, L50-33, and the revertant that emerged from the L50-33 strain were inoculated intranasally (3 x 10^5^ TCID_50_) into five-week-old male Syrian hamsters. The average of weight change is plotted and error bars represent SDs.

## STAR□Methods

### KEY RESOURCE TABLE

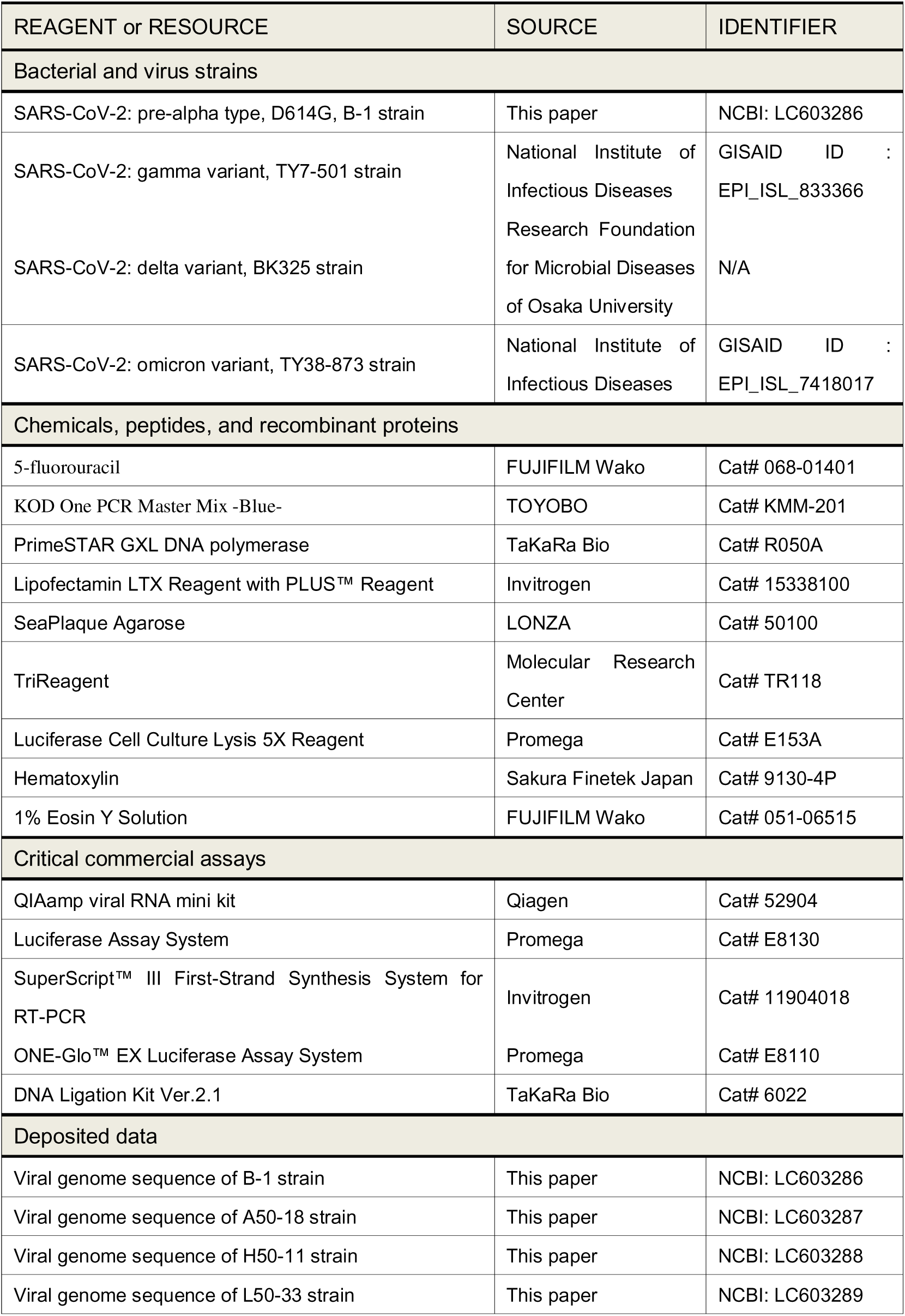
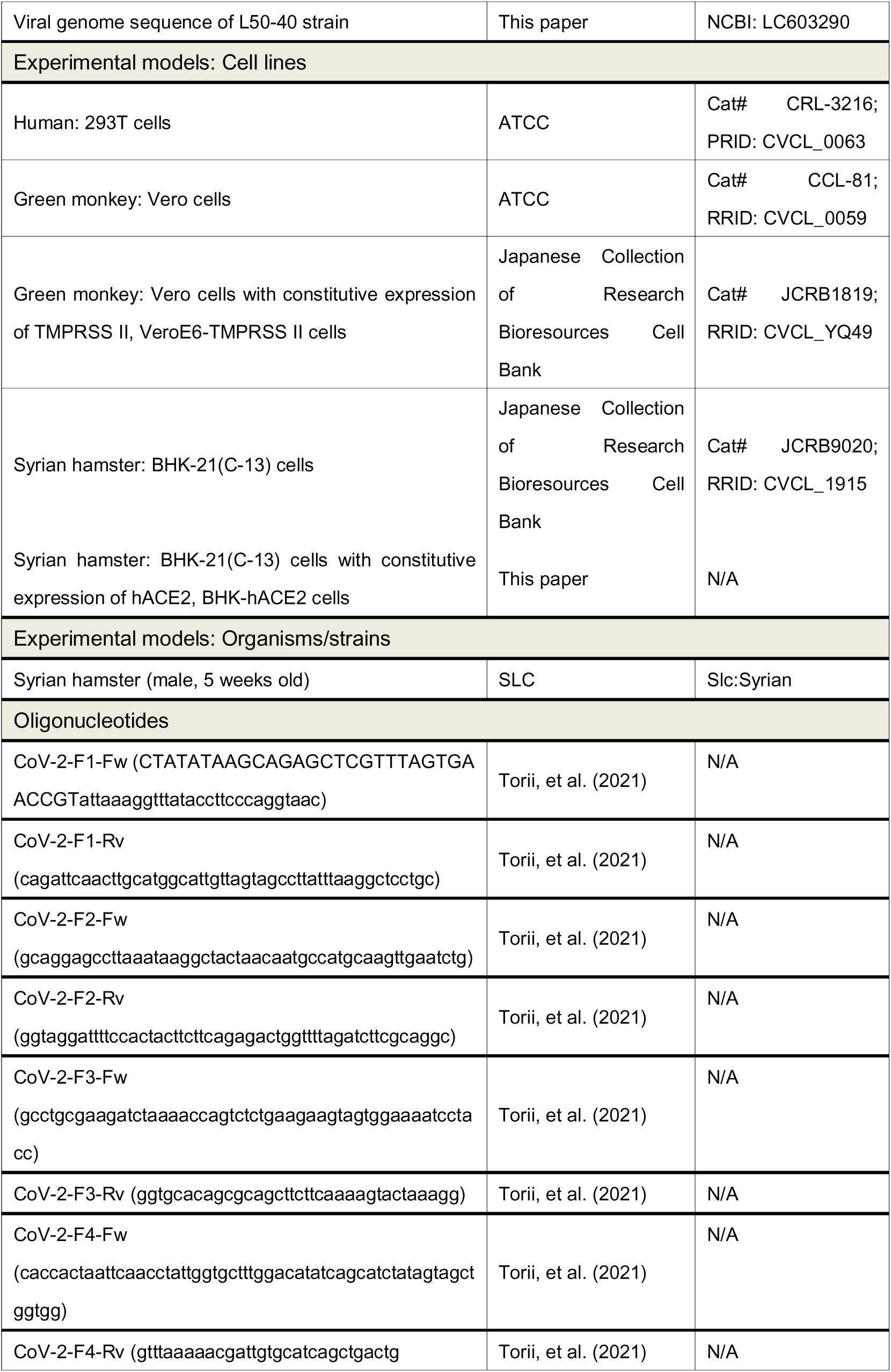
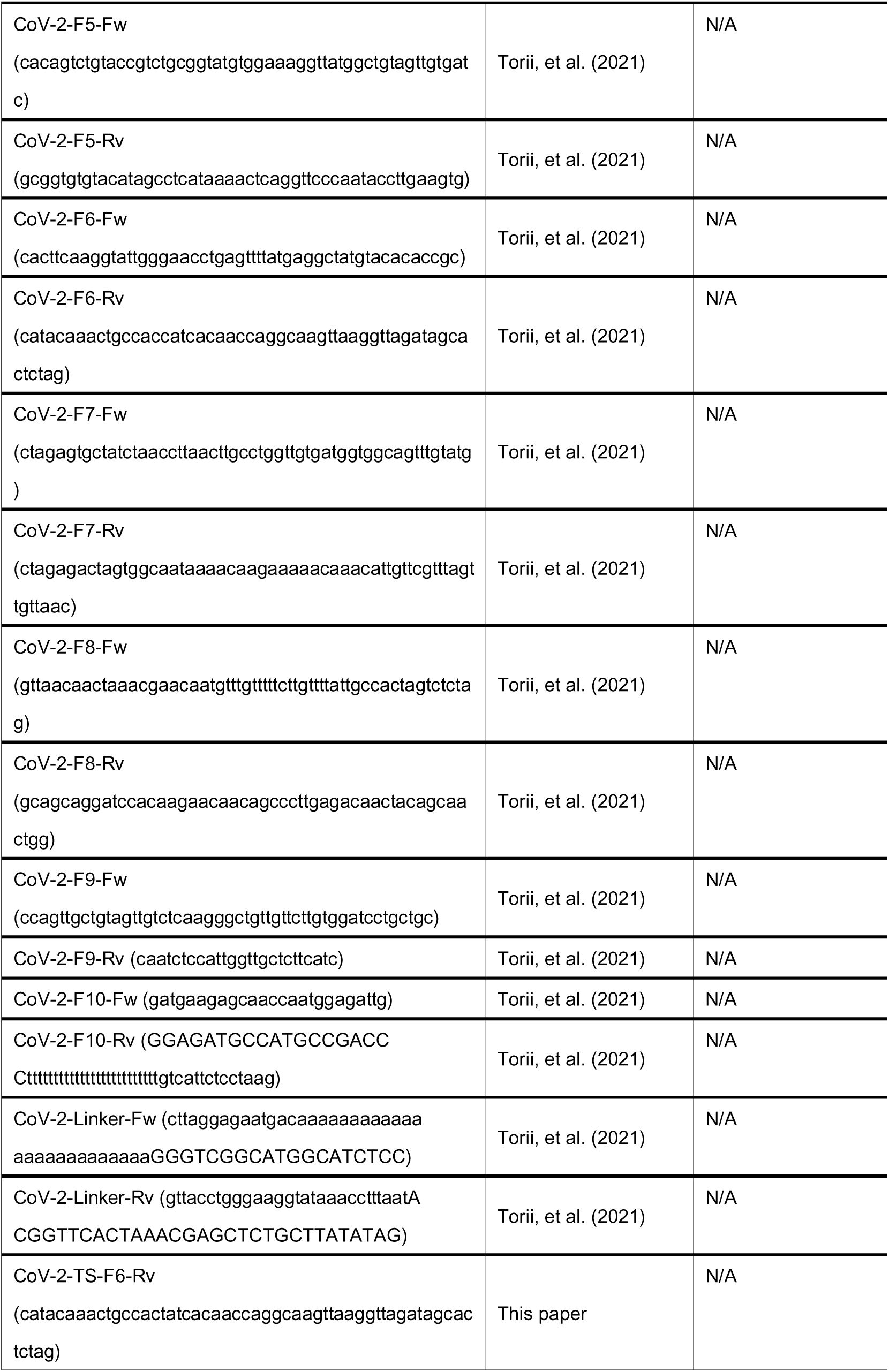
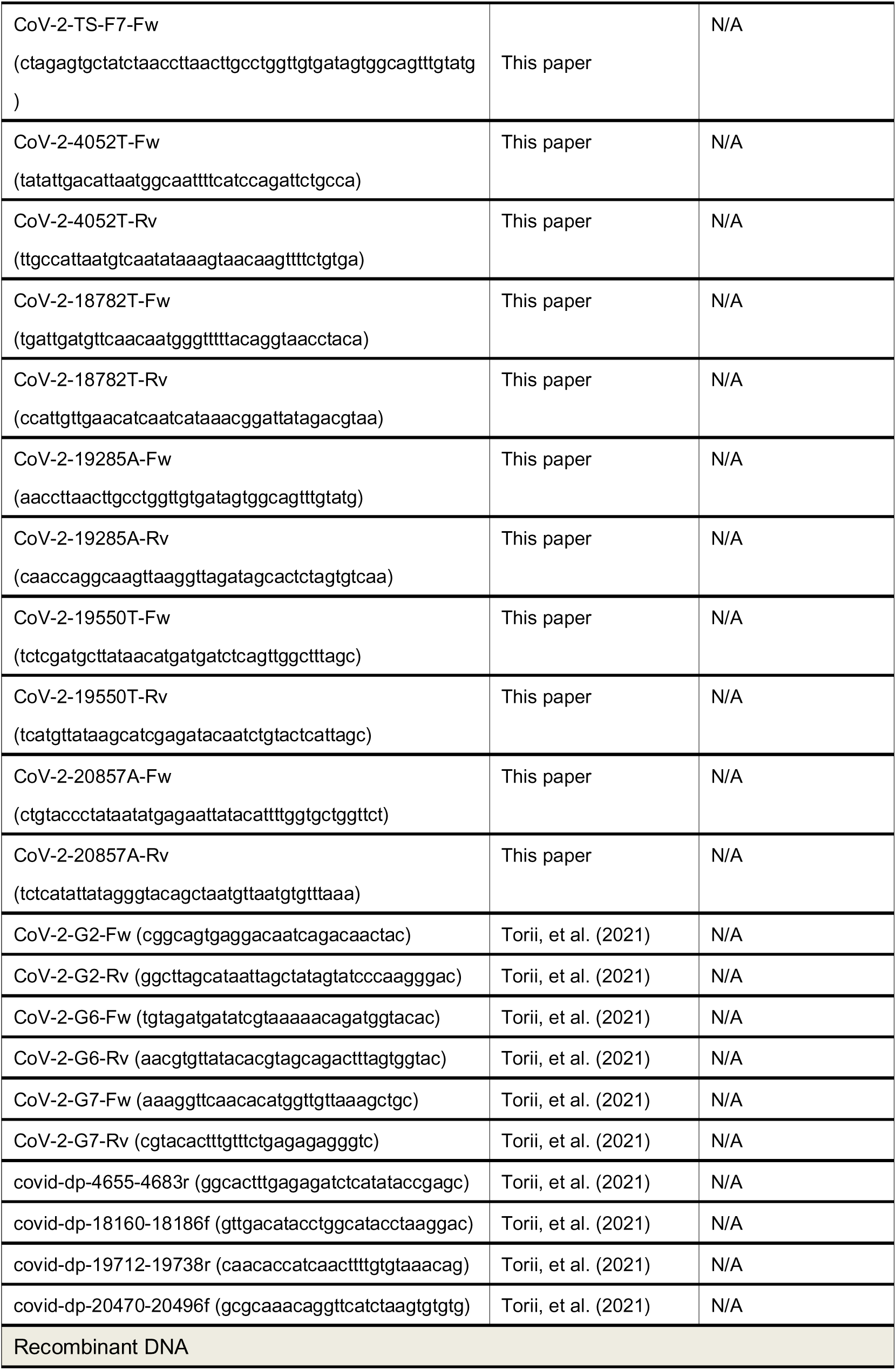
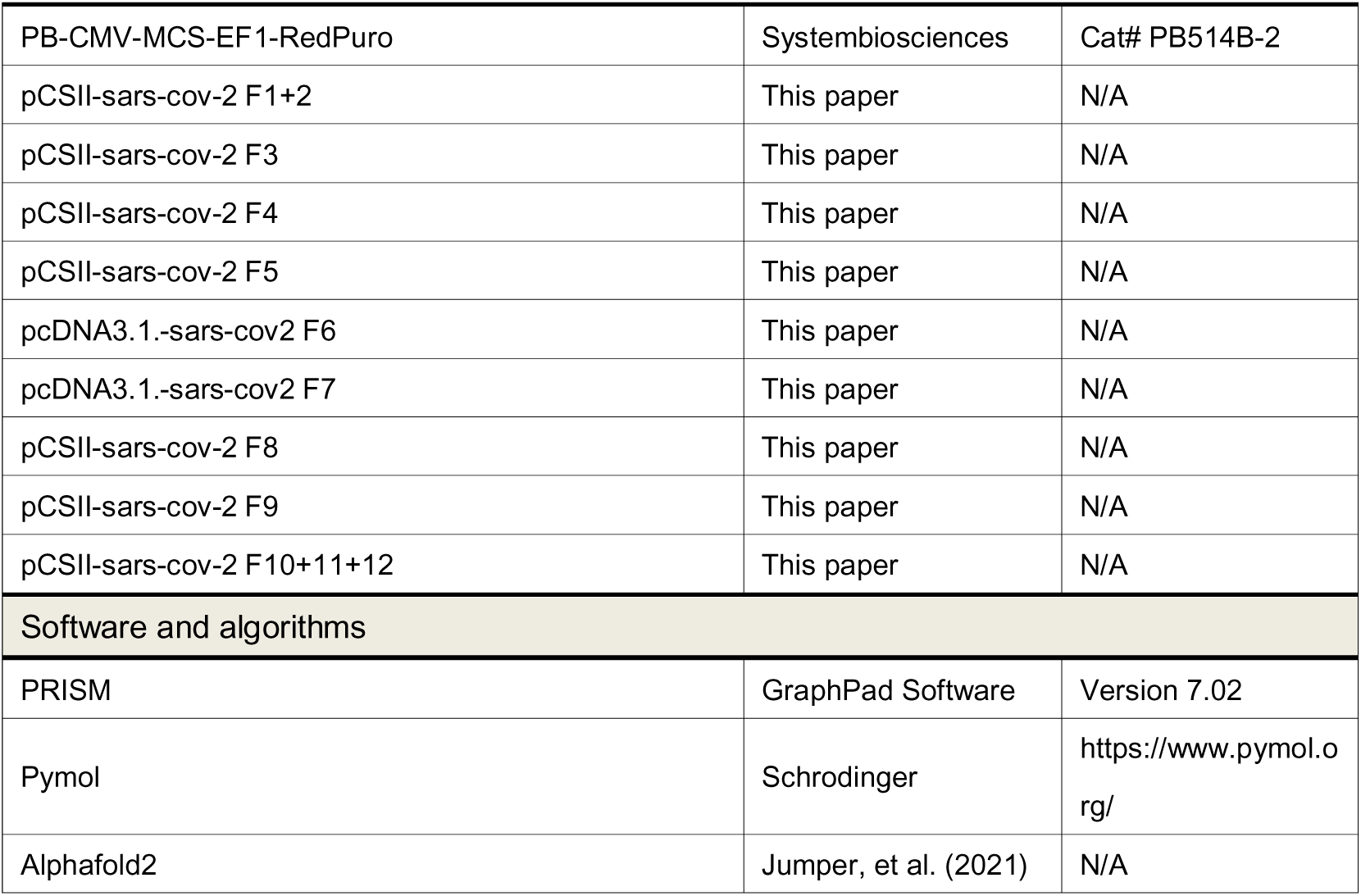

